# Defective subviral particles modify ecological equilibria and enhance viral coexistence

**DOI:** 10.1101/2022.05.03.490396

**Authors:** Adriana Lucía-Sanz, Jacobo Aguirre, Aurora Fraile, Fernando García-Arenal, Susanna Manrubia

**Affiliations:** School of Biological Sciences, Georgia Institute of Technology, Atlanta, GA, USA; Centro de Astrobiología (CSIC-INTA), Madrid, Spain; Grupo Interdisciplinar de Sistemas Complejos (GISC), Madrid, Spain; Centro de Biotecnología y Genómica de Plantas (CBGP), UPM-INIA, Madrid, Spain; Escuela Técnica Superior de Ingeniería Agronómica, Alimentaria y de Biosistemas, Universidad Politécnica de Madrid, Madrid, Spain; Systems Biology Department, National Centre for Biotechnology, CSIC, Madrid, Spain

**Keywords:** viral dynamics model, cooperation, ecological competition, subviral agents, viral coexistence

## Abstract

Cooperation is a main driver of biological complexity at all levels. In the viral world, gene sharing among viral genomes, complementation between genomes or interactions within quasispecies are frequently observed. In this contribution, we explore the advantages that flexible associations between fully fledged viruses and subviral entities, such as virus satellites, might yield. We devise a mathematical model to compare different situations of competition between two viruses and to quantify how the association with a satellite qualitatively modifies dynamical equilibria. The relevant parameter is the invasion fitness of each virus or of the virus-satellite tandem, which in the model depends on the transmission rate of viruses and on their effect on host survival. While in a virus-virus competition one of the viruses becomes eventually extinct, an association with a satellite might change the outcome of the competition to favor the less competitive virus (regardless of whether it is the helper virus or not) or to allow for the stable coexistence of the two viruses and the satellite. We hypothesize that the latter scenario, in particular, constitutes a parsimonious evolutionary pathway towards more stable cooperative associations, such as bipartite viral forms.

## 1 INTRODUCTION

There are many instances in the Virosphere of associations of viruses with kin or with subviral entities (Elena et al., 2014). The involved agents propagate independently, and engage in transient associations that can be contingent or necessary regarding successful completion of the replication cycle. Virus satellites, for instance, are subviral particles that require the assistance of a specific helper virus for its replication and/or encapsidation (Kassanis, 1962), see Fig. 1. Their association with the helper virus is therefore necessary for the satellite, but contingent in principle for the virus (though some viruses also require the satellite for encapsidation). Bipartite viruses, in turn, have their genome fragmented into two pieces that encapsidate separately but are essential in the viral cycle, so their mutual cooperation is necessary Sicard et al. (2016); Lucía-Sanz and Manrubia (2017).

**Figure 1.**
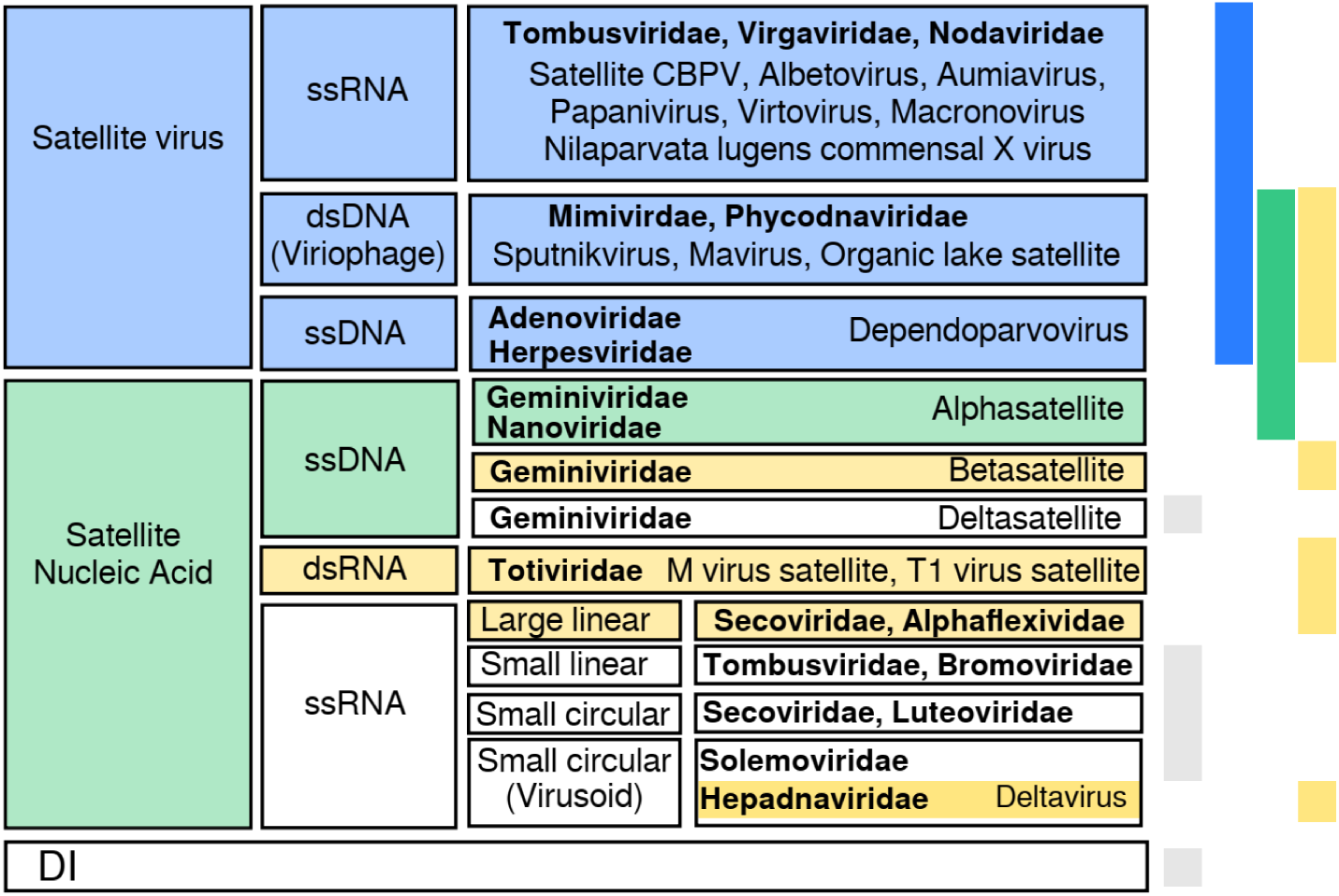
Summary of classes of satellite viruses. There is no straightforward classification of satellites, although distinguishable groups come to light when looking at the genetic material, helper virus family, host, or coded proteins. Here we show a possible classification into two groups, depending on whether they code for a capsid protein (satellite viruses) or not (satellite nucleic acid). Satellite viruses comprise plant satellites with jelly roll capsid proteins, dependoparvovirus and viriophages (Krupovic and Cvirkaite-Krupovic, 2012; Krupovic et al., 2016). Satellite nucleic acids include *α/β*–satellites and *Secoviridae* satellites coding for a replicase or a replication helper protein; M virus satellite expresses a toxin and HDV codes for an antigen. This group includes non-coding —circular— ssDNAs (Stanley et al., 1997) and RNAs with a compact folded structure with ribozyme activity (Hadid et al., 2017; Roossinck et al., 1992). Colors on the right bars show the protein each satellite codes for: capsid (blue), replicase (green), other (yellow). Grey stands for no coded protein. Helper virus families are shown in bold font; DI stands for defective interfering genomes (e.g., coated like defective interfering particles or uncoated).

Virus-satellite associations are ubiquitous in plants, and less frequent in other hosts. Examples infecting animals are the *Hepatitis δ virus* (HDV) (Makino et al., 1987), the genus *Dependoparvovirus* (Cotmore et al., 2014; Krupovic et al., 2016) that infects vertebrates, and virus-satellite associations that have bees (Ribière et al., 2010) and planthoppers (Nakashima et al., 2006) as hosts. There are also two cases of dsRNA satellites associated to the *Totiviridae* family that infect unicellular eukaryotes (Khoshnan and Alderete, 1995; Schmitt and Breining, 2002). In several cases, co-infection of animal hosts with the corresponding satellite results in an attenuation of viral symptoms (Nakashima et al., 2006; King et al., 2012; Yau et al., 2011).

Infections by multiple viruses and a variety of subviral entities are common in wild plants (Roossinck, 2005), opening up a high potential for mutual interaction. It has been observed that unrelated viruses within these mixtures do not seem to compete, but rather to cooperate, a fact that might explain their ubiquity (Leeks et al., 2018). Co-infection of a host with subviral particles usually modifies the pathogenicity of the helper virus (Roossinck, 2005; García-Arenal and Fraile, 2017). The spectrum of phenotypic modifications elicited in plants through such interactions is actually remarkably broad. Some viral species within the *Tombusviridae* family, for instance, take part in associations that include a variety of satellites coding for the capsid protein (Syller, 2003). *In vitro*, Tombusviruses spontaneously generate defective interfering particles (DIPs) that systematically lessen the symptoms of the infection by interfering with the replication process Havelda et al. (2005). However, co-infections with heterologous viruses, e.g. that of Potyvirus with Tombusvirus species as *machlomovirus of maize chlorotic mottle*, often increase the virulence of Tombusvirus (Scheets, 1998), as it also happens in other Potyvirus co-infections with heterologous viruses (Pruss et al., 1997). In Bromoviruses, virus-satellite associations can both attenuate or enhance the pathogenicity of the phenotype (Palukaitis and Roossinck, 1996). *Geminiviridae* is the family with the largest number of virus-satellite associations, often modifying the virulence and host-range of the virus (Rehman and Fauquet, 2009). Overall, while short-lived interactions may just involve an increase or decrease in viral accumulation levels (a common effect in infections where DIPs or satellites have spontaneously emerged), long-term interactions correlate with a variety of changes in transmission (Elena, 2011), host range (McLeish et al., 2019a), or cell tropism. Sustained interactions are in all likelihood a prerequisite to eventually develop a necessary, mutual interdependence between any two contingently interacting elements.

From an ecological perspective, phenotypic changes caused by transient associations with other viral and subviral agents affect viral dynamics, change viral epidemiology (Alcaide et al., 2020), and redefine the overall ecological role of the association (Betancourt et al., 2011; McLeish et al., 2019b; Syller, 2020), as compared to infections caused by a single virus. As such, they have to be subject to strong selection, since they may play a main role in virus survival by varying the cost that infection impinges on the host, changing adaptive strategies or, as we show in this contribution, qualitatively modifying the outcome of competition for hosts with co-circulating viral species. An integration of quantitative epidemiology at various levels, including in-host evolution and diverse adaptive strategies, is essential to better understand molecular associations and their ecological effects (Jeger, 2020).

Previous modelling studies have addressed the effect of co-infection by viruses and subviral particles unable to replicate on their own. Specific models have addressed the evolution of virulence in *Cucumber mosaic virus* over two hosts (Betancourt et al., 2013) and have shown that highly virulent phenotypes occur in mixed infections, in agreement with field observations. Dynamics of virus-DIPs have been also formally studied based on results of *in vitro* observations, revealing that dynamics in virus cultures can be oscillatory (Frank, 2000) but also intrinsically unpredictable under very general conditions (Kirkwood and Bangham, 1994); among others, DIPs can cause the extinction of the parental virus (Grande-Peérez et al., 2005). General models exploring properties of virus-satellite associations have often focused on conditions for coexistence of the partners, showing that it can be favored under structured demes (Szathmaéry, 1992) but is in general difficult under metapopulation dynamics (Nee, 2000). A general result for infections within a single host states that beneficial co-infection suffices to allow for the stable coexistence of different variants (Leeks et al., 2018). Still, the variety of models, and therefore mechanisms, identified for the stable coexistence of heterologous viral associations is reduced, probably as a result of our limited knowledge of the intricacies of such associations in nature, and to the small number of cases described (García-Arenal and Fraile, 2017). The state-of-the-art is similar in the case of necessary associations, such as in multipartite viruses, where specific mechanisms that may counteract the deleterious effects of independent propagation are still unclear (Lucía-Sanz et al., 2018; Zwart et al., 2021).

In this work, we present a simple model of virus-satellite association and evaluate its epidemiological effects when two monopartite viruses and a satellite that uses one of them as helper virus co-circulate in a host population. Our main result is that the virus-satellite association changes the possible stable equilibria and, in particular, promotes the long-term stable coexistence of both viruses and the satellite in a host population when, as most natural associations do, the satellite modulates the viral phenotype. In contrast, stable co-circulation of two monopartite viruses is not possible in the studied scenario, since one of the species unavoidable invades the host population in finite time.

## 2 MODELS

We address the epidemic propagation of viruses in a well-mixed host population through a set of differential equations representing the amount of susceptible hosts, *H*, of hosts infected by a virus, *X*, of hosts infected by the helper virus, *Y*, and of hosts simultaneously infected by the helper virus and the satellite, *S* (see Fig. 2). Capital letters, therefore, describe the abundance of hosts in each of four states: susceptible *H* or infected hosts in the infected states *X, Y*, and *S*; where needed, we will use lower-case letters to refer to virus *x*, helper virus *y*, and satellite *s*. The model, therefore, does not explicitly consider free viral populations. In our models, infections are persistent and there is no class of recovered hosts. This is the rule for plant viruses, but not necessarily so for animal viruses. Therefore, and although we generically speak of hosts all through the paper, our scenarios better apply to plant-infecting viruses and, as it has been defined, to any system where infections are persistent. Finally, note that we refer to competition for hosts in all instances, not to within-host competition.

**Figure 2.**
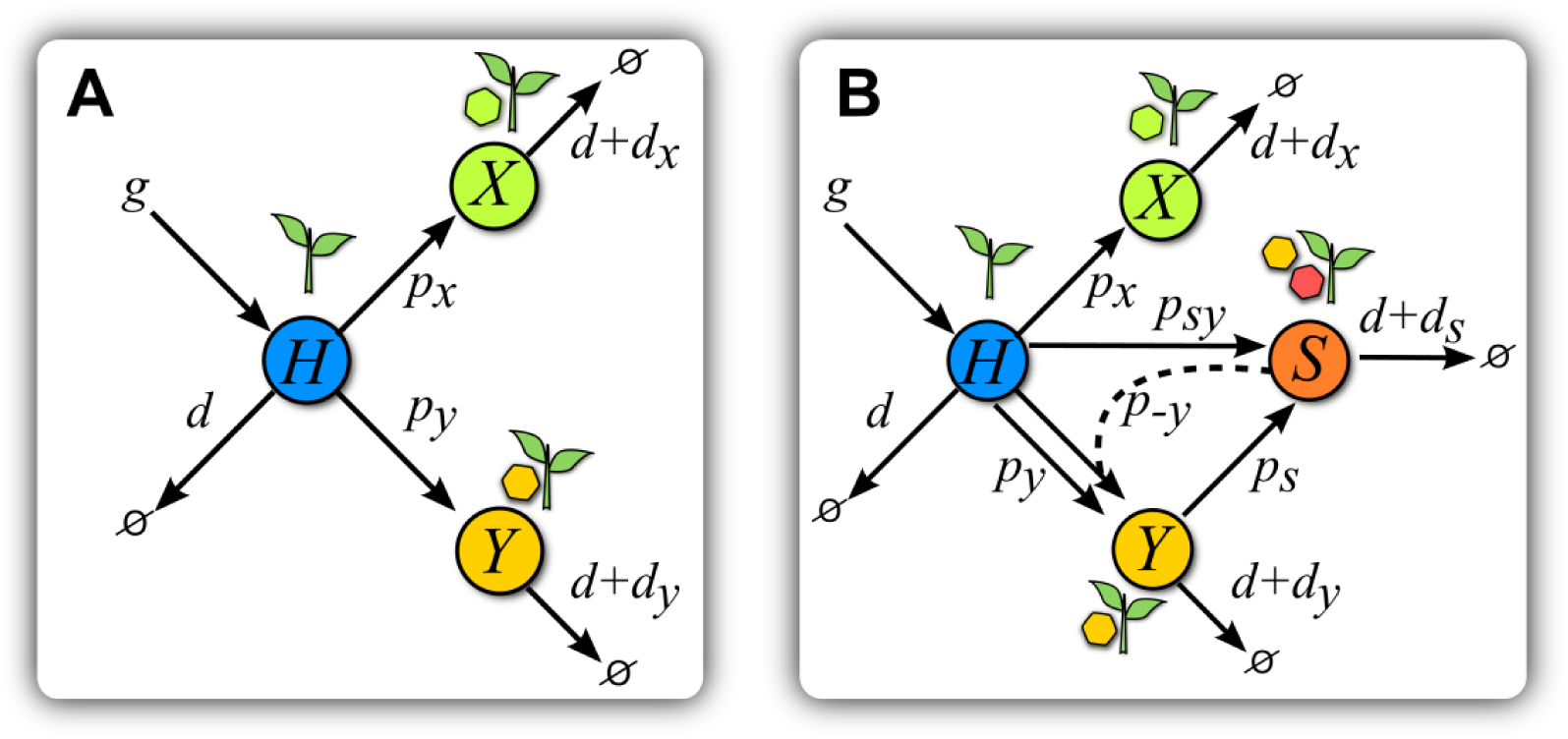
Scheme of models. (A) Scheme of Model V. Healthy hosts *H* are seeded at a constant rate *g* and die with basal rate *d*. Transitions to infected states happen at rates *p*_*x*_ and *p*_*y*_ for each virus. Infected hosts *X* and *Y* suffer from an increase *d*_*i*_ in basal mortality. (B) Scheme of Model S. The introduction of a satellite with *y* as helper virus adds new transitions with respect to (A). Hosts in class *Y* can get infected at a rate *p*_*s*_ by a satellite upon contact with hosts in class *S*. Hosts in this latter class are simultaneously infected by virus *y* and its satellite, and have an increase *d*_*s*_ in their mortality rate. Infection of *H* plants by the helper virus and the satellite simultaneously occurs at a rate *p*_*sy*_ upon contact with class *S*. A co-infected plant *S* can also infect *H* plants only with virus *y* at a rate *p*−*y*. The color code is maintained all through the paper.

We keep the model intendedly simple so as to derive general principles arising from competition for hosts between the association of a helper virus and a satellite and a second monopartite virus. We do so to emphasize that relevant ecological dynamics do depend on the success of a specific strategy in front of alternative others. In order to highlight the main processes involved in such competition, we have made some simplifying assumptions. For instance, we assume superinfection exclusion between the two viruses, such that hosts cannot be simultaneously infected by the two monopartite viruses. This is a convenient simplification to obtain exact results; as a consequence, coexistence in the context of our model means “ecological coexistence”, in the sense that the two viral strains and the satellite stably co-circulate in the host population under conditions that will be made explicit later. A second simplification is that of a mean-field approximation (common in paradigmatic epidemic models (Hethcote, 2000)), where the dynamics are described through differential equations and space is not explicitly modeled, therefore assuming that hosts interact homogeneously through averaged values.

### 2.1 Competition between two viruses: Model V

The general dynamics of the process are as follows. Healthy hosts appear at a constant rate *g* and decay or die at a natural rate *d*. The amount of healthy, susceptible hosts at time *t* is *H(t)* and that of hosts infected by either virus is *X(t)* and *Y(t)*. If no confusion arises, we will obviate the explicit dependence on *t*. Contacts between susceptible and infected hosts cause infection of susceptible hosts at rates *p*_*x*_ and *p*_*y*_, respectively, that stand for the transmission rates of either virus. Infection by *x* or *y* increases the death rate of the host in amounts *d*_*x*_ and *d*_*y*_. This scenario corresponds to a simple situation where two fully competent viruses compete for hosts (see Fig. 2A), and is described by the following equations:

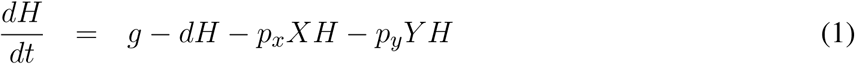

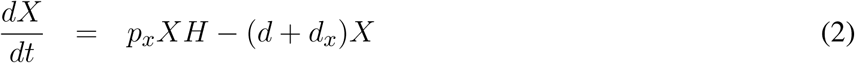

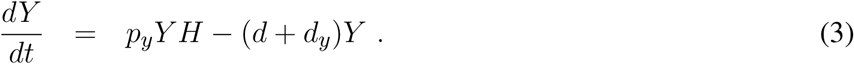

In the absence of viruses (*X* = 0, *Y* = 0), the abundance of hosts stabilizes at *H*^*^ = *g/d*, as obtained by solving *dH/dt* = 0. Therefore, Eq. 1 entails a maximum carrying capacity of hosts in the absence of parasites. From now on, and given its relevance in determining the existence and stability of the various solutions, as it will be shown, the quantity *d/g* will be called host turnover.

It is a consequence of the symmetry between eqs. (2) and (3) that one of the viruses always displaces the other —with rare marginal exceptions, as we show in the results. Alternatively, for a small enough host replacement, both viruses become extinct: infected hosts die before new susceptible hosts are available to maintain the epidemic.

### 2.2 Competition between a virus and a helper virus with a satellite: Model S

In the former scenario of competition between two viruses, we are now introducing a possible association with a satellite that can only multiply in presence of its helper virus. Due to the symmetry between the equations describing the dynamics of each virus, Eqs. (2) and (3), the choice of the helper virus does not affect the results, so we selected virus *y* without loss of generality. Class *S*, corresponding to hosts infected by the tandem *y* − *s*, is affected by an increase *d*_*s*_ in mortality.

Due to the dependence of the satellite on its helper virus for replication, *s* can only be transmitted (to classes *H* or *Y*) through contacts with class *S*, either to healthy hosts, at rate *p*_*sy*_, or to hosts already infected with *y*, at rate *p*_*s*_. In agreement with our superinfection exclusion, hosts in class *X* cannot be infected by classes *Y* or *S*. Finally, hosts in class *S* can also transmit only the helper virus under contacts with class *H*, this process occurring at rate *p*−*y*. A scheme of the interactions of this model can be seen in Fig. 2B, and the set of equations considering all of the previous processes reads:

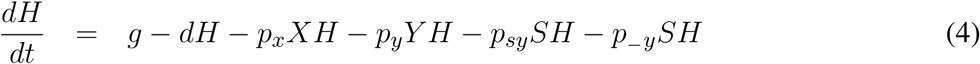

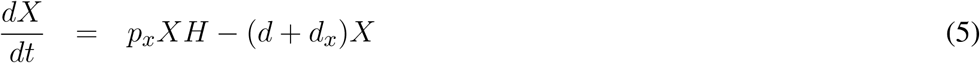

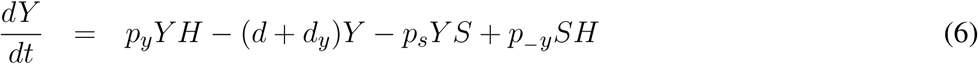

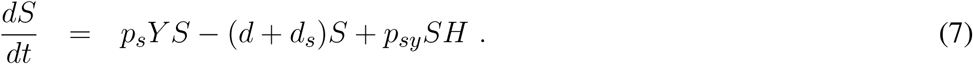

Note that, if we set *S*(*t* = 0) = 0 as an initial condition, this system is in practice equivalent to Model V. This nonetheless, the formal inclusion of the satellite modifies the study of the fixed points and stability of the system, as we show in the Supplementary Information.

The satellite non-trivially breaks the symmetry between the dynamics of hosts in classes *X* and *Y*, and therefore between the two corresponding viruses. As we will show through analytical and computational analyses, the satellite can benefit or impair its helper virus (thus impairing or benefiting virus *x*), or can promote the stable coexistence of all parts.

## 3 RESULTS

In order to clarify the effects of the introduction of a satellite in the outcome of competition between two monopartite viruses infecting the same host (model S), we begin by analyzing the stable solutions of the dynamics in the absence of the satellite (model V). All parameters in the models (death and transmission rates and host growth) are positive defined. The results obtained for model V are not conceptually new, but serve to properly compare the dynamics of the more complex model S with a baseline situation. Fixed points of the dynamics are the solutions to 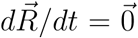, where 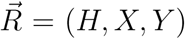 (model V) or 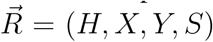 (model S). The values of the variables satisfying the fixed point equations are denoted *H*^*^, *X*^*^, *Y* ^*^, and *S*^*^ in the following. These values have to be equal to or larger than zero for the solution to have biological meaning: this results into conditions for existence and non-negativity that the model parameters have to fulfill. The study of stability of the solutions has been carried out using standard methodology of dynamical systems, as detailed in the Supplementary Information. Stability of the solutions also yields conditions on formal relationships that the model parameters have to satisfy. In the next two sections we summarize the solutions of models V and S and their stability properties, together with their ecological interpretation.

### 3.1 Coexistence is unstable in a system of two viruses in competition for a host population

Depending on model parameters, model V has four different, positive and stable solutions: i) none of the virus is able to invade the population of hosts and both get extinct; ii) and iii) one of the two viruses stably coexists with the host and the other virus becomes extinct; iv) the two viruses coexist in the population. These four solutions map on three main extended regions in the space of parameters.

An important quantity arising in the formal analysis of the model is the ratio between the transmission rate of a virus, *p*_*i*_, and its overall effect on host death rate, *d* +*d*_*i*_, where *i* = *x, y*. This ratio is an instance of the *invasion fitness F*_*i*_ = *p*_*i*_*/*(*d* + *d*_*i*_) of each virus, and determines which strategy is uninvadable (Lehmann et al., 2016). The ratio *d/g* is a measure of the replacement of healthy hosts. Smaller *d/g* means shorter times are needed for the appearance of susceptible healthy hosts. Invasion fitness *F*_*i*_ and host turnover *d/g* are the two main quantities determining the relevant equilibria of the system. A summary of such solutions and their stability is reported in Table 1 (see Supplementary Material for further details). Figure 3 shows a representative example of the localization in parameter space of the four possible solutions for model V. In that particular example, we have fixed *d* = 0.05, *d*_*y*_ = 0.25, *p*_*y*_ = 0.1, and *p*_*x*_ = 0.1 and explored the {*g, d*_*x*_} plane. Changes in parameter values shift the boundaries between different solutions but do not cause qualitative changes. We observe large regions where both viruses become extinct or where either *x* or *y* coexist with the host.

**Table 1.**
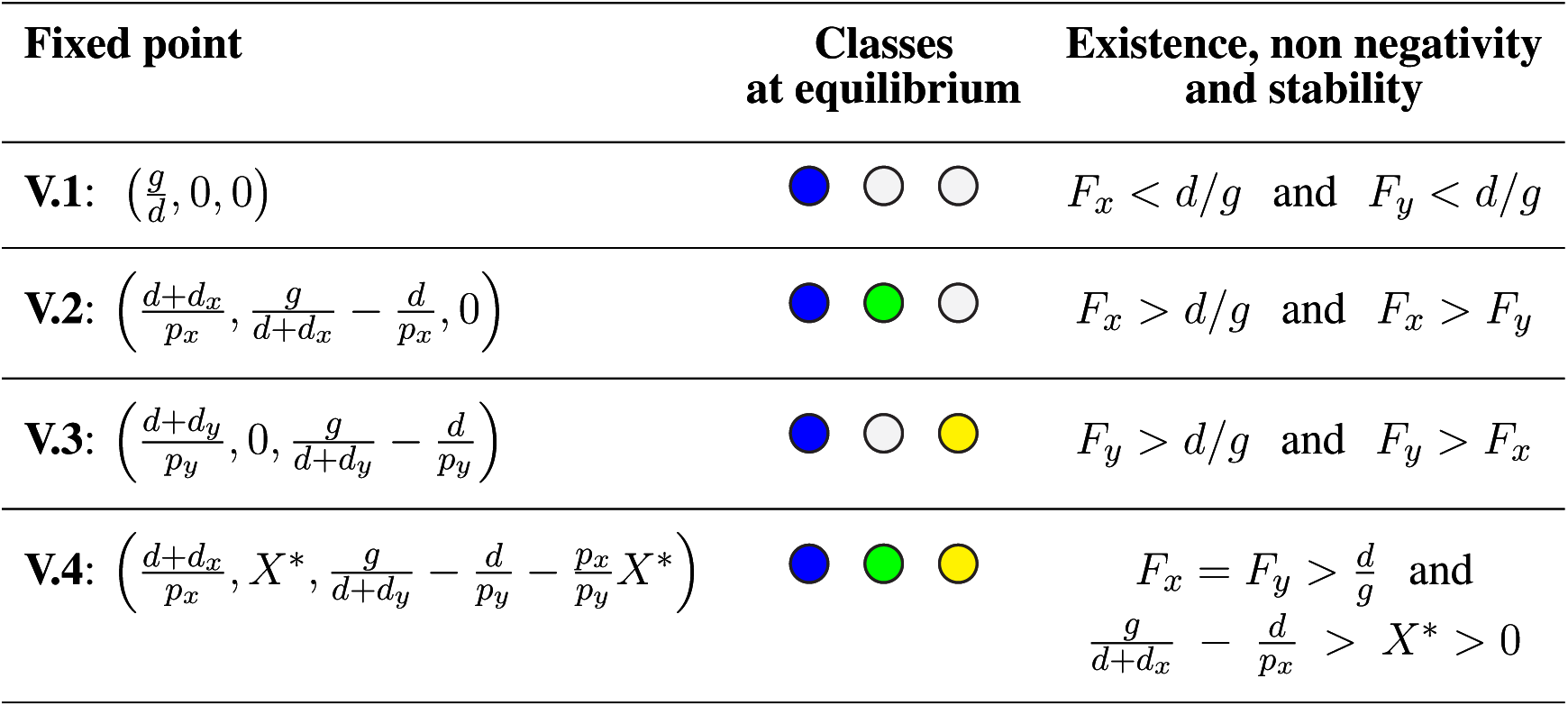
Properties of the solutions of model V. The left column shows the analytical solution (fixed point of the dynamics, (*H*^*^, *X*^*^, *Y* ^*^)); the central column symbolically illustrates the states with positive population at equilibrium (colored circles), gray circles are for classes with zero population. The right column specifies the conditions for existence, non-negativity and stability of each solution.

**Figure 3.**
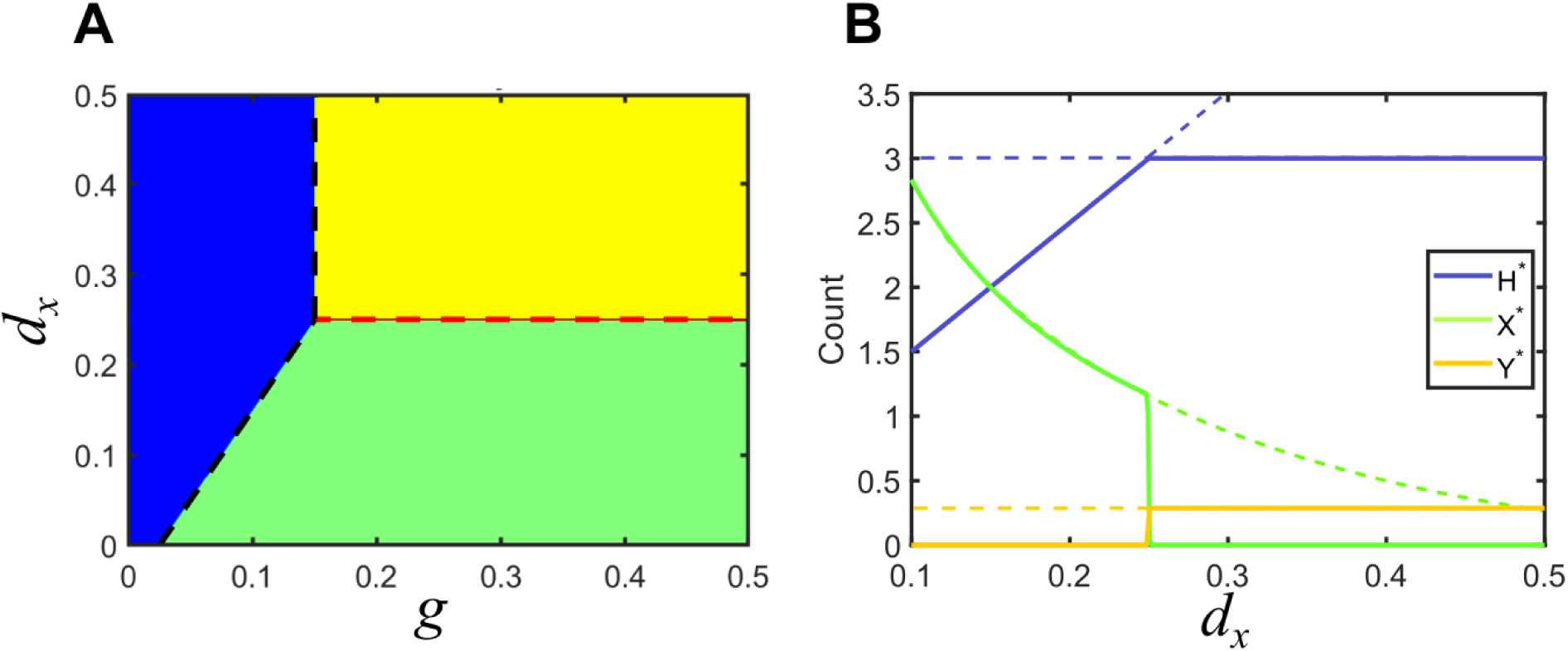
Solutions for Model V in parameter space. (A) Phase space of solutions in the plane {*g, d*_*x*_ }. In the blue region, none of the viruses can invade the population of hosts, *d/g* > *F*_*i*_, *i* = *x, y*. The black dashed line indicates the boundary where *d/g* = *F*_*i*_. Green and yellow regions correspond to solutions that satisfy *F*_*x*_ > *F*_*y*_ and *F*_*y*_ > *F*_*x*_, respectively, where one of the viruses invades the population of hosts and displaces the other. The red dashed line corresponds to the marginal coexistence solution. (B) Bifurcation diagram for a cross section of the phase space in A corresponding to *g* = 0.5. Lines indicate the amount of each class at the fixed point, as shown in the legend. Dashed lines represent unstable solutions, while solid lines correspond to stable solutions. Green and yellow regions in A correspond to *d*_*x*_ *<* 0.25 and *d*_*x*_ > 0.25. Vertical lines show the degenerate solution of coexistence of the two viruses for *d*_*x*_ = 0.25. Note that the values of *H*^*^ and *Y* ^*^ become independent of *d*_*x*_ in the region where *y* is the invading virus (and thus *x* becomes extinct). Parameters used: *d* = 0.05, *p*_*x*_ = 0.1, *p*_*y*_ = 0.1, *d*_*y*_ = 0.25.

#### 3.1.1 Extinction of both viral populations

If host turnover is larger than invasion fitness for both viruses (solution **V.1**), the viral populations becomes extinct. This region corresponds to the blue area in Fig. 3A, and describes a solution where the rate of appearance of new susceptible hosts is slower than the time needed by any of the viruses to invade the population: in absence of sufficient susceptible hosts, any parasite becomes extinct.

#### 3.1.2 Survival of one viral population

This situation holds whenever the invasion fitness of the two viruses is different and at least one of them is higher than host turnover: *F*_*i*_ > *F*_*j*_ and *F*_*i*_ > *d/g* for at least one *i, j* ∈ *{x, y }*. The solution is symmetric under the exchange *i* ↔ *j*. These situations correspond to solutions **V.2** and **V.3**, represented as green and yellow regions in **Figure 3.A**. The abundances of susceptible and infected hosts depends on all model parameters.

#### 3.1.3 Coexistence of both viruses

This is a marginal solution sitting at the boundary separating the two solutions where one of the viruses invades and the other becomes extinct. Coexistence only occurs when the two viruses have identical invasion fitness, and both are higher than host turnover, *F*_*y*_ = *F*_*x*_ > *d/g*. This condition maps to a single line in parameter space. Further, since the equilibrium values of *X*^*^ and *Y* ^*^ are degenerated, a second condition establishes that none of them be zero (in which case the solution would correspond to cases **V.2** or **V.3**). This means that, at this boundary, the amount of hosts infected by *x* or *y* is not unique, as illustrated by the vertical line in Fig. 3B, where the amounts of *X* and *Y* appear superimposed. The degeneration of the solution is also reflected in its analytical form **V.4**, since *Y* ^*^ is given as a function of *X*^*^ (see Table 1 and Supplementary Information).

It is not to be expected that any natural system strictly meets the conditions for coexistence here derived. In the example in Fig. 3, any minor inaccuracy in the value of *d*_*x*_ would kick the system out of coexistence and cause the extinction of one of the viruses. In general, arbitrary changes in *d*_*x*_, *d*_*y*_, *p*_*x*_ or *p*_*y*_ would cause the system to depart from the (quasi)stable solution. Therefore, in more realistic situations, stochasticity, noise, or heterogeneity of any kind would push the system away from this marginal solution and into one of the extended regions where either one or both viruses become extinct. Therefore, co-circulating different viral species are, in general, not to be found in a host population under the conditions of model V.

### 3.2 Dynamics with a co-infecting satellite

The introduction of the satellite breaks the symmetry of equations describing the dynamics of the two viruses. In this new situation, it is not straightforward to predict which of the viruses or associations is going to succeed in invading the host population. A variety of outcomes is possible depending on the new parameters.

We first extend our definition of invasion fitness to the virus-satellite association,

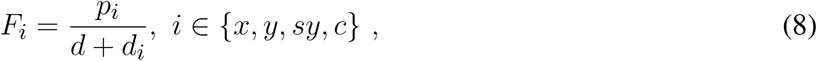

where for convenience we define *p*_*c*_ = *p*_*sy*_ + *p*−*y* (with *d*_*c*_ = *d*_*sy*_ = *d*_*s*_). Note that, since the likelihood of independent transmission is larger than joint transmission, *F*_*c*_ > *F*_*sy*_.

The system of equations (4-7) has seven different fixed point solutions (for all parameters taking positive values), summarized in Table 2. Four of them coincide with the solutions for model V, namely: extinction of the two viruses (**S.1**), invasion of virus *x* (**S.2**) or virus *y* (**S.3**) and (marginal) coexistence of hosts with viruses *x* and *y* (**S.4**). Three new solutions appear where the satellite co-circulates with one or two of the viruses. The first of them (**S.5**) corresponds to the extinction of virus *x* and the stable coexistence of the tandem virus *y*-satellite with the host. In two more solutions (embraced under **S.6**) the three agents (virus *x*, virus *y* and satellite) stably co-circulate in the population of hosts. In the following, we examine in detail solutions **S.5** and **S.6** to discuss how the introduction of the satellite modifies the different equilibria depending on the phenotypic effects of its association with the helper virus.

**Table 2.**
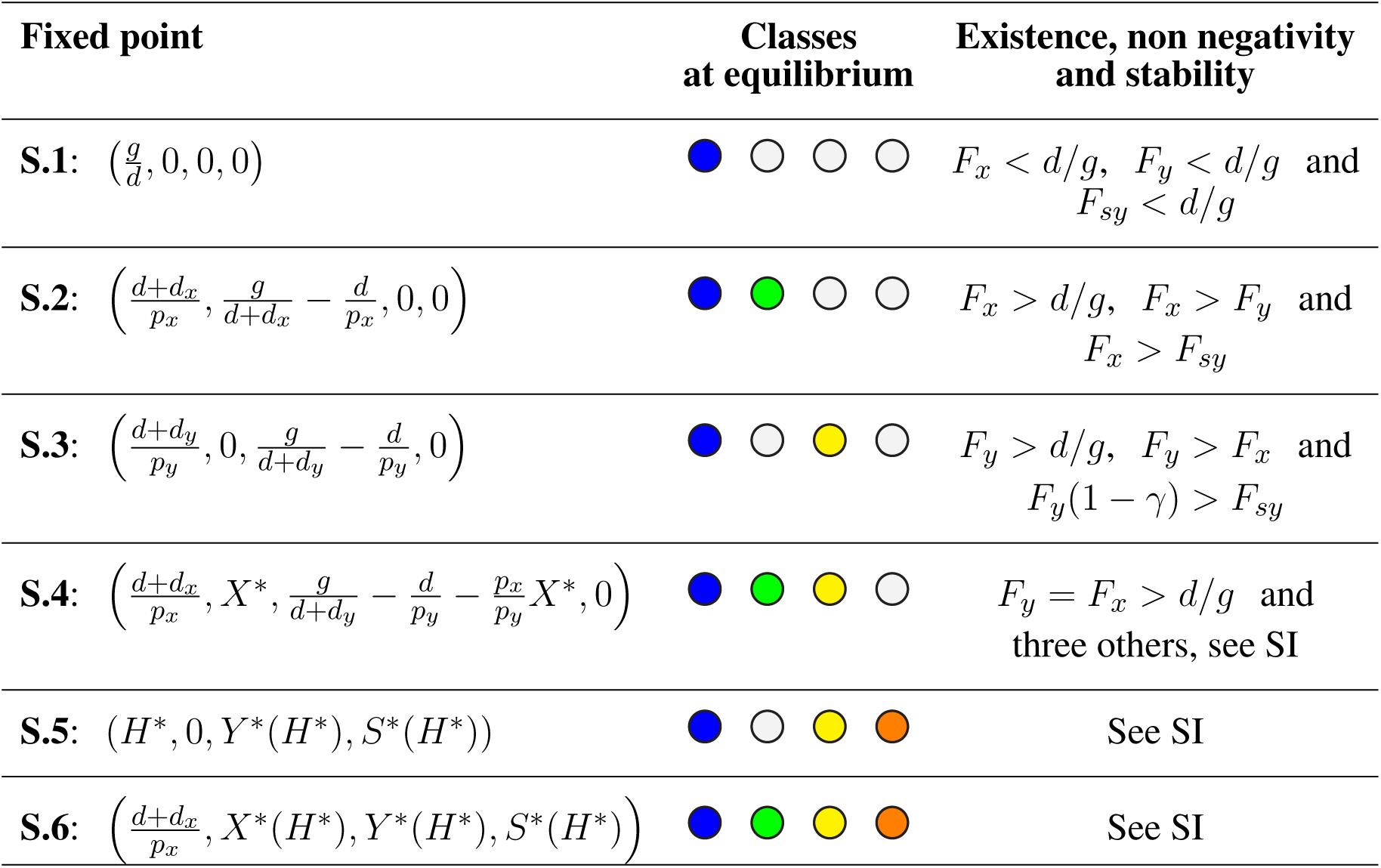
Properties of the solutions of model S. The left column shows the analytical solution (fixed point of the dynamics, (*H*^*^, *X*^*^, *Y* ^*^, *S*^*^)); the right column summarizes all conditions for existence, non-negativity and stability of that solution. We have defined *γ* = (*g/*(*d* + *d*_*y*_) − *d/p*_*y*_)*p*_*s*_*/*(*d* + *d*_*s*_), see main text. Solution **S.4** is degenerate in the same sense than **V.4** was: there is a continuum of values of *X*^*^ and *Y* ^*^ that satisfy the fixed point equations. Further, it is a marginally stable solution, since parameter values have to be fine-tuned for it to persist. This is the reason it requires strict conditions that we detail in the Supplementary Information. Coexistence solution **S.6** takes two different forms, therefore embracing two different fixed points (see main text and SI for further details).

#### 3.2.1 Neutral association with a satellite does not modify the outcome of competition

If the satellite behaves as a commensal parasite, co-infection with its helper virus does neither modify the regions of different equilibria nor their stability. Formally, this occurs when *F*_*c*_ = *F*_*y*_. Fig. 4 depicts the regions for the different equilibria and shows that the region corresponding to solution **V.3** is now split into two subregions embracing solutions **S.3** and **S.5**: in the former, the satellite becomes extinct; in the latter, it co-circulates with virus *y*.

**Figure 4.**
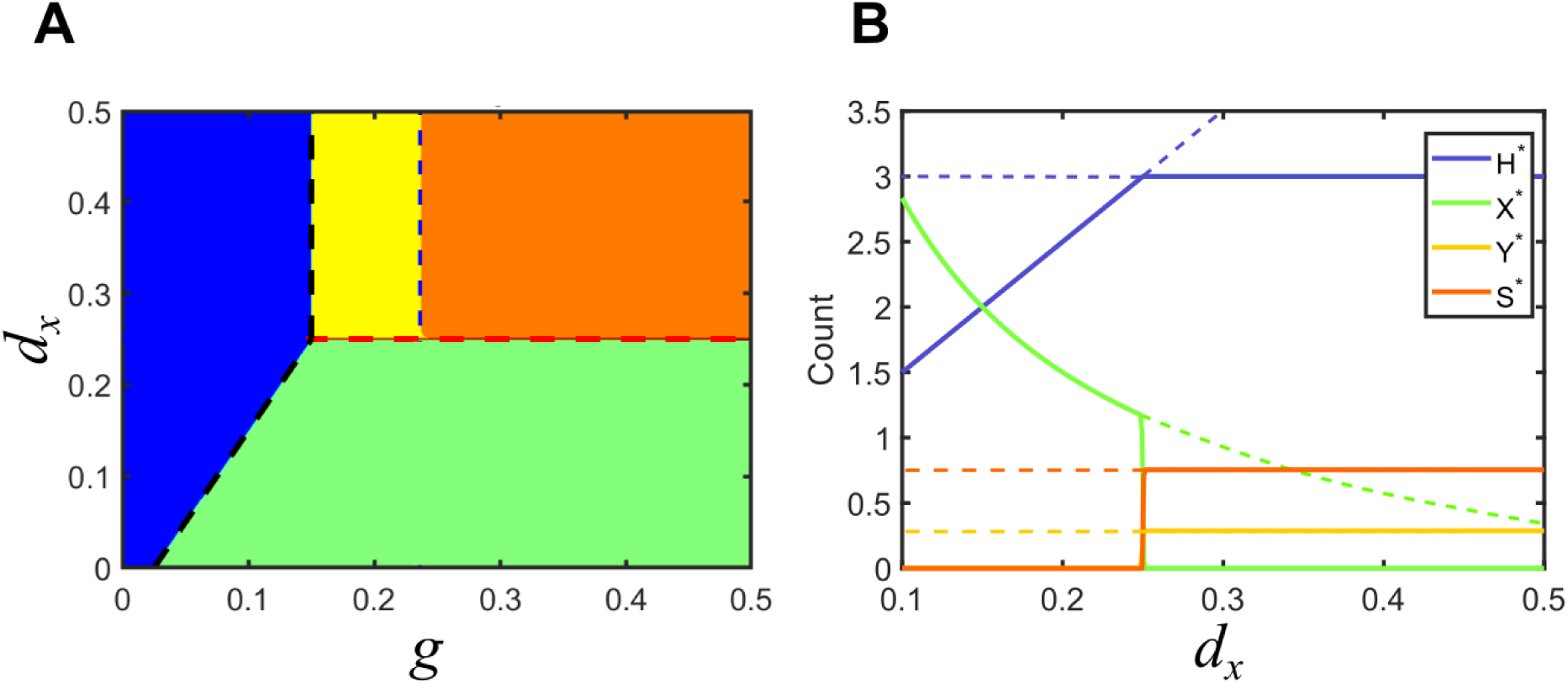
Solutions for model S in parameter space: commensal satellite. (A) Phase space of solutions in the plane {*g, d*_*x*_} for a commensal satellite (*F*_*c*_ = *F*_*y*_). The description of regions corresponding to solutions **S.1** (in blue, viruses cannot infect hosts), **S.2** (in green, virus *x* invades and virus *y* becomes extinct) and **S.3** (in yellow, virus *y* invades and virus *x* becomes extinct) are analogous to those described for Model V. In the new orange region, the satellite stably coexists with its helper virus, **S.5**. Red and black dashed lines are as in Fig. 3. The blue dashed line signals the boundary separating the regions with and without co-circulating satellite. Note that the absence of virus *y* prevents the stable circulation of the satellite (below *d*_*x*_ = 0.25). (B) Bifurcation diagram of a cross section of A, for *g* = 0.5. As above, lines stand for the amount of each class at equilibrium. Dashed lines correspond to unstable solutions, while solid lines correspond to stable solutions. Solutions collide with the degenerated solution **S.4** at *d*_*x*_ = 0.25. Parameters used: *d* = 0.05, *p*_*x*_ = 0.1, *p*_*y*_ = 0.1, *d*_*y*_ = 0.25, *p*_*s*_ = 0.8, *p*_*sy*_ = 0.04, *d*_*s*_ = 0.3, *p*_−*y*_ = 0.0766.

Invasion of virus *y* without the satellite (**S.3**) holds when *F*_*y*_(1 − *γ*) > *F*_*sy*_, where *γ* = (*g/*(*d* + *d*_*y*_) − *d/p*_*y*_)*p*_*s*_*/*(*d* + *d*_*s*_). This condition is shown as a vertical, blue dashed line in Fig. 4. As the invasion fitness *F*_*sy*_ of the virus-satellite association increases, it becomes harder for the helper virus to survive on its own until, eventually, solution **S.5** becomes stable and substitutes **S.3**: in **S.5**, *Y* ^*^ ≠ 0 and, simultaneously, *S*^*^ ≠ 0. In general, this means that satellites are persistent in helper virus populations, but the virus still infects a subpopulation of hosts in the absence of the satellite. The size of this region depends on the ability of virus *y* to infect susceptible hosts alone when jumping from joint virus-satellite infections. Therefore, it progressively narrows if *p*−*y* decreases. A successful strategy from the viewpoint of the satellite, therefore, could be to evolve towards associations that make the *y*-virus more dependent on *s* for transmission, progressively selecting increases in *p*_*s*_ that could eventually turn *p*−*y* negligible. That is, the association would evolve from commensal to mutualistic.

#### 3.2.2 A mutualistic satellite can prevent extinction of the helper virus

Consider the situation in model V where *F*_*x*_ > *F*_*y*_, with *F*_*x*_ > *d/g* (solution **V.2**): virus *y* is at a disadvantage and therefore unable to invade the population of hosts. In that situation, cooperation with a satellite can prevent its extinction if the satellite provides a sufficiently high increase in invasion fitness to its association with *y*. This occurs when the invasion fitness of the virus-satellite association is slightly larger than *F*_*x*_. A region of parameters that was previously occupied only by solution **S.2** becomes now bistable, such that depending on the initial conditions it can lead to the elimination of *y* (and the concomitant disappearance of the satellite), or to the extinction of *x* in favor of the virus-satellite association (solution **S.5**), see Fig. 5.

**Figure 5.**
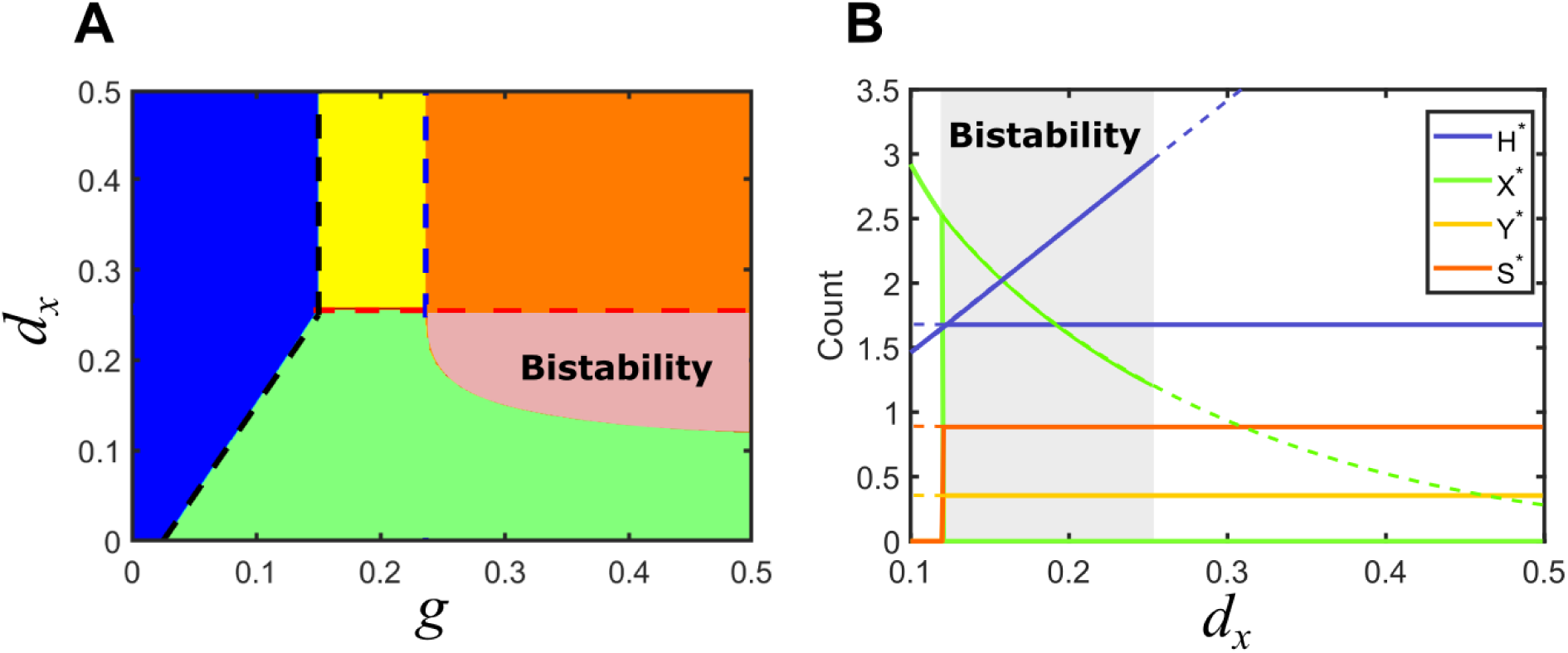
Solutions for model S in parameter space: mutualistic satellite. (A) Phase space of solutions in the plane {*g, d*_*x*_} for a mutualistic satellite. Here, *F*_*c*_ > *F*_*y*_. Some of the previous equilibrium solutions are maintained, as long as the inequalities that define them remain unchanged. However, a region of bistability appears, where solutions **S.2** and **S.5** are possible for the same parameters, depending on the initial conditions. (B) Bifurcation diagram of a cross section of A, for *g* = 0.5. As above, lines stand for the amount of each class at equilibrium. Dashed lines correspond to unstable solutions, while solid lines correspond to stable solutions. A region of bistability for 0.12 *< d*_*x*_ *<* 0.25 is shown in gray. A bifurcation where two solutions appear occurs at *d*_*x*_ = 0.12; solutions collide with the degenerated solution **S.4** and stability is modified at *d*_*x*_ = 0.25. Parameters used: *d* = 0.05, *p*_*x*_ = 0.1, *p*_*y*_ = 0.1, *d*_*y*_ = 0.25, *p*_*s*_ = 0.8, *p*_*sy*_ = 0.04, *d*_*s*_ = 0.3, *p*_−*y*_ = 0.2.

At the boundary that separates the region of bistability (where both **S.2** and **S.5** are possible) from solution **S.2** as a unique solution, the values of *H*^*^ for each solution coincide and equal *H*^*^ = (*d* + *d*_*x*_)*/p*_*x*_, as Fig. 5B shows. By using the explicit expression for *H*^*^ obtained for **S.5** (see Supplementary Information) one can obtain an implicit analytical expression for the lower bistability boundary.

From an evolutionary viewpoint, satellites that confer their helper virus an increase in invasion fitness may be a low-cost solution to guarantee rapid adaptation to new hosts, since such an adaptive strategy, in principle, does not require adaptive mutations.

#### 3.2.3 A parasitic satellite can promote viral coexistence

Consider now the situation in model V where *F*_*y*_ > *F*_*x*_, with *F*_*y*_ > *d/g* (solution **V.3**): virus *x* is at a disadvantage and virus *y* has invaded the host population. An association of *y* with a parasitic satellite can be a burden to the helper virus and diminish its relative advantage while, at the same time, it could potentially benefit *x*, the virus that was initially eliminated in the competition between the two monopartite viruses. It turns out that the association with a satellite under the conditions above does not necessarily entail, in contrast to previous equilibria, the elimination of either *x* or the *y*-satellite tandem. Actually, if the invasion fitness of the association virus *y*-satellite is such that *F*_*c*_ *< F*_*x*_ *< F*_*y*_, then virus *x* can simultaneously coexist with virus *y* and the satellite in regions where it became extinct when the satellite was absent. The conditions above correspond to the relevant solution embraced under **S.6**, that we label **S.6.a**, see SI. There is a second degenerate solution that requires *F*_*c*_ = *F*_*x*_ = *F*_*y*_ (solution **S.6.b**) that has not biological relevance, in the same sense as discussed above for other degenerate solutions.

Figure 6 illustrates the parameter space for the case of a parasitic satellite and shows the appearance of an extended region of coexistence. The boundary limiting the coexistence region can be analytically calculated by noting that, at that boundary, the values of *X*^*^ for solutions **S.5** and **S.6.a** coincide and are equal to zero. See Supplementary Information for an explicit expression of the latter solution.

**Figure 6.**
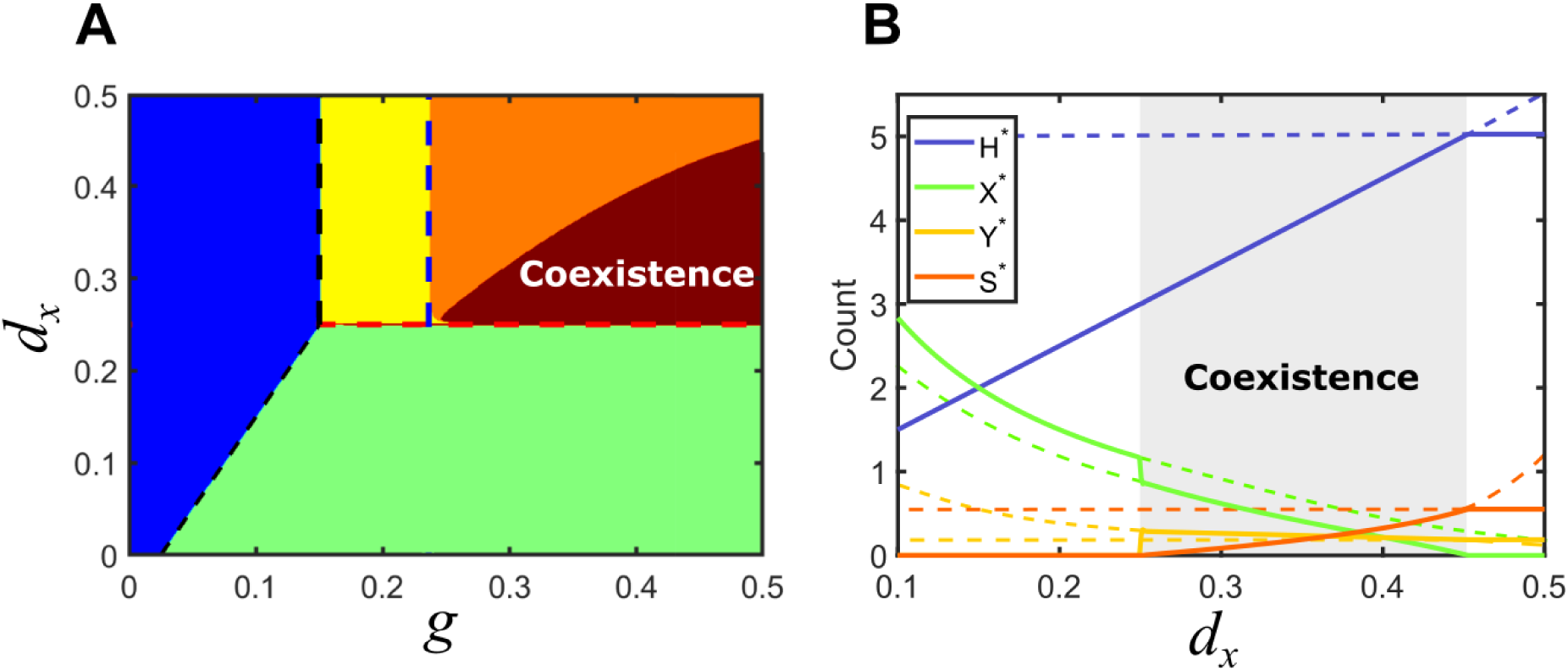
Solutions for model S in parameter space: parasitic satellite. (A) Phase space of solutions in the plane {*g, d*_*x*_} for a parasitic satellite in the particular case *F*_*c*_ *< F*_*y*_. A region of co-existence occupies part of the area where the virus *y*-satellite eliminated virus *x* if the satellite was commensal or mutualistic. (B) Bifurcation diagram of a cross-section of A, for *g* = 0.5. A region of co-existence of all the populations is shown in gray for 0.25 *< d*_*x*_ *<* 0.4529, corresponding to solution **S.6**. Solutions collide with the degenerated solution **S.4** at *d*_*x*_ = 0.25; a bifurcation occurs at *d*_*x*_ = 0.4529. Parameters used: *d* = 0.05, *p*_*x*_ = 0.1, *p*_*y*_ = 0.1, *d*_*y*_ = 0.25, *p*_*s*_ = 0.8, *p*_*sy*_ = 0.04, *d*_*s*_ = 0.3, *p*_−*y*_ = 0.016.

The case of coexistence is an interesting and ecologically relevant situation where the association with a parasitic satellite is only partly detrimental for virus *y*, while it actually benefits both virus *x* and the diversity of circulating viral species (and subviral entities).

### 3.3 Increased dependence between helper virus and satellite

We have shown in the previous sections how the asymmetric relationship between virus *y* and the satellite (the virus can propagate and replicate on its own, while the satellite depends on the helper virus) breaks the symmetry of the simple model V in a non-trivial way. This causes in particular the appearance of regions of bistability and coexistence that depend on the phenotype of the association, as compared to that of virus *y* alone.

There are two other situations implicit in model S and in the solutions analyzed so far that are worth discussing: the case where the helper virus cannot propagate if the host is not co-infected with the satellite, and the limit case where the “helper virus” needs the “satellite” both for propagation and replication. The former case (section 3.3.1) still distinguishes between the two categories of helper virus and satellite, while in the latter (section 3.3.2) the association has turned necessary: the two partners that can no longer be identified as helper virus and satellite, since their interdependence is mutual and alike. Actually, these two cases might represent intermediate steps in a possible parsimonious evolutionary pathway towards bipartitism.

### 3.3.1 A satellite virus coding for genes essential in transmission

Consider a situation where virus *y* can replicate but not propagate without the concurrence of the satellite. Formally, this situation is implemented by setting *p*_*y*_ = 0, meaning that hosts in class Y are not infective. This scenario agrees with various common natural situations (see Fig. 1) where a satellite virus codes for the capsid needed for the encapsidation of the helper virus or is essential for vector transmission. Examples of the latter are the requirement of *groundnut rosette satellite* for the transmission of *groundnut rosette virus*, of RNA4 for transmission of *beet necrotic yellow vein virus* (Palukaitis et al., 2008) or the dependence of umbraviruses on luteoviruses for transmission (Taliansky and Robinson, 2003).

In our formal context, this restriction implies that only hosts in class S can generate virions with virus *y* able to infect other susceptible hosts. This is the only way in which the Y class is fed, since the term *p*_*y*_*Y H*, corresponding to infections caused by contacts between Y and H hosts, is now absent. Taking an evolutionary perspective, the association between a subviral particle with multiplicative ability and a satellite that endows it with propagative ability appears beneficial for the particle. From the viewpoint of the satellite, it might be also convenient, in order to ensure survival, to carry a highly-valued genome piece, as a capsid or others involved in transmission, that can be likely used by a variety of replicative particles. From this perspective, evolutionary associations between replicative entities and satellites coding for essential genes could be highly dynamic, successful adaptive strategies.

### 3.3.2 The helper virus can neither replicate nor propagate without the satellite

The subsequent case corresponds to a situation where the partners become mutually dependent both for multiplication and propagation. The formal representation of this situation corresponds to setting *Y* = 0, since the “helper” virus cannot replicate in the absence of the “satellite”, and therefore class Y is empty —in the same sense that a class infected only by satellites is irrelevant for the dynamics, and therefore dismissed in our models from scratch. When the set of Eqs. (4)-(7) is reduced to this case, one can see that they are equivalent to model V under the correspondence *Y* → *S, d*_*y*_→ *d*_*s*_, *p*_*sy*_ → *p*_*y*_, and *p*−*y* = 0, since class *Y* is absent by definition. Still, there is an important difference due to the fact that virus *y* and the satellite do not necessarily propagate as a unit, but as two independent particles. As a result, *p*_*sy*_ in the equivalent formulation may take low effective values, meaning that the probability of simultaneous infection by two independently propagating particles is always lower than the probability of infection by a single propagating particle. Therefore, this situation better describes a bipartite virus, where the genome is divided into two segments that are independently transmitted, where both fragments are needed to successfully complete the viral cycle, and where there is a cost due to independent propagation.

## 4 DISCUSSION

We have introduced a simple epidemic model to analyze stable states in a system formed by two fully fledged viruses and an associated satellite. This model contains the minimum set of players with symmetric (between the two viruses) and asymmetric interactions (between a virus and a satellite) of different nature (commensal, parasitic or mutualistic) able to display a range of ecological equilibria when competing for a common host. Note that our results only refer to long-time equilibria (i.e. to stable coexistence in a mathematical sense), and therefore do not apply to transient coexistence. In natural systems, the time needed for one viral species to cause the elimination of a second one (corresponding, for example, to solutions **V.2** or **V.3**) can be so large that transient co-circulation of various viral species could be the most common situation: this transient coexistence should not be mistaken by solution **V.4**, which strictly refers to stable, long-term coexistence. A detailed analysis of how long it takes to attain ecological equilibrium and a quantification of typical evolutionary times for the emergence of new viral species infecting a given host would be needed, among others, to discriminate between transient and actual coexistence. Such an analysis is out of the scope of the present contribution, but it is important to keep in mind that, due to the long times involved in ecological dynamics, natural systems can be out-of-equilibrium most of the time.

In the light of our results, it is worth discussing the major assumption we have made, that is superinfection exclusion. Our second assumption, the use of a deterministic description of the dynamics, mostly affects the stability of marginal solutions (i.e., those solutions demanding a precise coincidence of certain parameters). The incorporation of stochasticity of any kind (in the form of space, which entails probabilistic contagion, finite populations, heterogeneous contacts, noise or any form of disorder), as it would occur in any natural system, turn solutions **V.4, S.4**, and **S.6.b** irrelevant, since they become unstable under any minor modification of most parameter values.

Assuming superinfection exclusion is useful to obtain exact analytical results, and therefore to gain insight into major ecological effects of virus-satellite associations. If this simplification is lifted, further interaction terms between viruses *x* and *y*, between *x* and the satellite, or even among the three entities, and new equations accounting for different states with various combinations of co-infecting entities have to be considered. Both the number of equations involved and their complexity would increase, preventing the achievement of exact expressions for equilibria and their stability, and therefore masking the interpretation of generic results. This nonetheless, and since we are arguing that coexistence of different viruses and subviral particles in the same host population might be a stepping stone towards stable mutually dependent associations (as bipartitism), let us be more explicit about this point and sketch how the relaxation of that assumption could affect the results here obtained. The simplest scenario corresponds to commensal co-infection, in which case the qualitative results here presented remain unchanged: propagation of different entities in the host population could be independently analyzed, and the state of each individual host at a given time would be the product of probabilities of being infected with any of the viruses or sets of partners considered in the model. If co-infection causes changes in phenotype, however, the nature of the different equilibria might change, but such changes would be qualitatively analogous to these explored here.

Take the following example as illustration. Suppose we include in our equations hosts co-infected with viruses *x* and *y*, and let us call *Z* this new class. The dynamical equation for *Z* would read

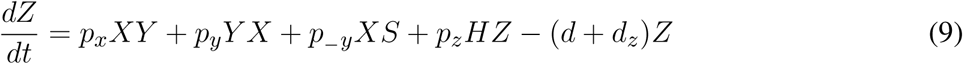

The terms in the right-hand side of the equation indicate, from left to right, the rate of infection of hosts in class *Y* under a contact with class *X* (virus *x* is transmitted), of hosts in class *X* under a contact with *Y* (virus *y* is transmitted), of hosts in class *X* under a contact with *S* when only *y* is transmitted, new infections of susceptible hosts under contacts with *Z* when both viruses are simultaneously transmitted (this introduces the new rate *p*_*z*_) and, finally, the modification of the death rate due to the co-infection with both viruses (new rate *d*_*z*_). Equations for *dH/dt, dX/dt* and *dY/dt* would also acquire new terms coming from the new transitions to class *Z* (equation for *dS/dt* would not be modified since there are no possible transitions between *S* and *Z*, since the latter does not contain the satellite). Note that this enlarged model Z would have all fixed points we have described for model S (when *Z*^*^ ≠ 0) and some new ones corresponding to *Z*^*^ = 0. The analysis would proceed along similar lines, and the actual challenge would then be to characterize the new conditions under which coexistence occurs, since these could take much more involved mathematical expressions. Still, the mechanisms underlying coexistence, in particular, would not differ from the ones identified in this study as long as interaction terms take mathematical forms analogous to those considered here, that is, if interactions occur in pairs. This is not a limiting assumption, since all general contagion processes involve only pair contacts, regardless of whether multiple particles of various types are transmitted through the contact. For example, if another class *W* infected by *x, y* and *s* is considered, it can be easily shown that its dynamical equation only involves pair interactions again, even if the three entities can be simultaneously transmitted (for example through contacts between *H* and *W*). We therefore conclude that the basic model explored in this work contains essential mechanisms for coexistence resulting from asymmetrical interactions among the involved partners, and its results can be conceptually extrapolated to scenarios with a larger number of players.

This problem of identifying coexistence conditions and the involved partners also admits a complementary viewpoint. Instead of considering an enlarged dynamical system with many interacting viral and subviral entities, and struggle to identify the conditions ensuring coexistence, we could take a more pragmatic approach and ask which is the subset of viruses and subviral agents able to coexist when pooled from a large ensemble of viruses and subviral particles that infect a given host. This subset would be, by definition, an ensemble fulfilling the conditions for coexistence in that host, and it could vary from host to host. From this viewpoint, the result of evolving towards regions of coexistence —should this be a situation under selection— is equivalent to drawing the subset of viral species and subviral entities able to coexist. This latter scenario conceptually agrees with common observations of multiple viral species (and possible subviral particles) persistently co-infecting the same host, especially in plants.

Indeed, some of the equilibrium solutions and the overall scenarios we have found in our models and their extensions find their counterpart in natural situations, and do share some of their difficulties. While co-infection by multiple viral species or strains is common, detection and identification of involved parties and the nature of their interactions presents major problems in etiological and epidemiological studies (Jeger, 2020). In our modelling approach, we can just describe regions of parameter space where coexistence is or is not possible, but only a quantification of interactions and parameters can allow for a more specific identification of partners and their interactions. For example, in our solution **S.6.a**, we show that due to synergy between virus *y* and the satellite, *x* can stably infect the host, while it became extinct in the absence of *s*. This would be the case of the obligatory mutualism between *Pea enation mosaic virus 1* (PEMV1) and *Pea enation mosaic virus 2* (PEMV2), where PEMV1 provides a capsid required for PEMV2 transmission while PEMV2 provides a movement protein required for systemic movement of PEMV within the infected plant (Doumayrou et al., 2016).

Other studies have shown that the level of satellite abundance in virus-satellite associations is highly variable, with up to 100% in some regions for helper virus population in *Rice yellow mottle virus* satRNA (Pinel et al., 2003) to very low, as in *Cucumber mosaic virus* satRNA (Palukaitis and García-Arenal, 2003). These observations are compatible with solutions **S.5** and **S.6.a**, where the level of co-infected hosts in class *S* depends on several different model parameters. In particular, it grows when the rate of joint *y*-*s* transmission, *p*_*s*_ increases, and *vice versa*, reaching 100% in situations where, in the context of our model, both the replication and the propagation of *y* become fully dependent on the presence of *s*. Still, we cannot tell the specific mechanism (i.e. parameter value, or set of), allowing for coexistence.

Our main qualitative result is that stable coexistence of all players is possible under a broad range of parameters. Though biological and epidemiological outcomes of interactions among co-infecting viruses seem to be hardly predictable (Syller, 2020), long-lived loose interactions as those arising between the agents in our model create favorable circumstances for evolution towards more permanent associations and to promote recombination between dissimilar viral and subviral particles (Elena, 2011; García-Arenal et al., 2003). For example, recombination has led to the emergence of new species, both in DNA viruses as begomoviruses (Chakraborty et al., 2008), and in RNA viruses, as is the case of *Watermelon mosaic virus*, which arose through recombination of two legume-infecting viruses (Desbiez et al., 2011).

All examples of satellite viruses summarized in Fig. 1 represent necessary, asymmetric associations that have likely evolved from more loose interactions between helper viruses and satellites. In a parsimonious evolutionary scenario, it is sensible to assume that a monopartite virus coding for all functions needed to complete the viral cycle modified at some point its genome through the association with a satellite coding for genes providing new or different functions, either improving its packaging efficiency or its transmissibility, or modifying its infecting phenotype. A colonization of a new host where the genes provided by the satellite was instrumental could have also been a plausible evolutionary pathway, followed by the elimination of the old, now useless, equivalent gene in the viral genome. This sort of reasoning agrees with hypotheses relating high viral diversity to advantages at early phases of host colonization (Gonzalez-Jara et al., 2009), where loose associations could quickly modify the viral phenotype without the need of incorporating genetic changes (Lucía-Sanz and Manrubia, 2017).

Coexistence of viral and subviral agents in a single host may act as well as stepping stones towards mutually dependent necessary associations, as bipartite viral forms. Symmetric associations characterized by mandatory complementation between the parties could arise through different evolutionary pathways. They could evolve from asymmetric associations, as these described in the previous paragraph, when the survival of the helper virus becomes more dependent on the satellite. Also, they could be generated *de novo* through associations between complementary, defective particles. Finally, fully fledged viruses of dissimilar origin could produce a new phenotype and, eventually, streamline their genomes to minimize functional redundancy, in a sort of viral Black Queen association (Morris et al., 2012). The three pathways above could eventually yield what is usually described as a bipartite virus. Further particles could be added to the system through similar processes, causing in a parsimonious way the emergence of multipartite viral forms.

## AUTHOR CONTRIBUTIONS

AL, JA and SM participated in the overall conception and design of the study; SM designed the models; AL implemented the models and performed the simulations; AL and JA carried out the analytical calculations. AF and FG made substantial contributions to the discussion and interpretation of the results. AL and SM wrote the manuscript. All authors contributed to manuscript revision, read, and approved the submitted version.

## SUPPLEMENTAL DATA

A Supplementary Information text file includes details on calculations retrieving the fixed points of the dynamics and conditions for stability.

## DATA AVAILABILITY STATEMENT

The computational code required to generate all the results in this study is publicly available at https://github.com/aluciasanz/satcompet.git

