## Supplementary Information for "Defective subviral particles modify ecological equilibria and enhance viral coexistence"

### 1 ANALYTICAL SOLUTIONS AND STABILITY ANALYSIS OF MODEL V

The stationary solutions —or fixed points— $(H^*, X^*, Y^*)$ of model V are obtained by equating to zero the ordinary differential equations describing the model ($dH/dt = 0$, $dX/dt = 0$ and $dY/dt = 0$) and solving for the values of the variables, $H^*$, $X^*$ and $Y^*$:

$$0 = g - dH^* - p_x X^* H^* - p_y Y^* H^* \tag{1}$$
$$0 = p_x X^* H^* - (d + d_x) X^* \tag{2}$$
$$0 = p_y Y^* H^* - (d + d_y) Y^* \ . \tag{3}$$

Positive and stable solutions of the system correspond to the asymptotic state of the variables of the system, for sufficiently long time. Solutions are stable when the system returns to that stationary state if its variables are slightly perturbed. For the solution to be biologically meaningful, some conditions for existence and non-negativity of all variables have to be fulfilled, regardless of whether the fixed point is stable or unstable. Four stable non-negative equilibria solutions are found when Eqs. (1–3) are solved,

$$\begin{aligned}
\textbf{V.1: } (H^*, X^*, Y^*) &= \left(\frac{g}{d}, 0, 0\right) \\
\textbf{V.2: } (H^*, X^*, Y^*) &= \left(\frac{d+d_x}{p_x}, \frac{g}{d+d_x} - \frac{d}{p_x}, 0\right) \\
\textbf{V.3: } (H^*, X^*, Y^*) &= \left(\frac{d+d_y}{p_y}, 0, \frac{g}{d+d_y} - \frac{d}{p_y}\right) \\
\textbf{V.4: } (H^*, X^*, Y^*) &= \left(\frac{d+d_x}{p_x}, X^*, \frac{g}{d+d_y} - \frac{d}{p_y} - \frac{p_x}{p_y}X^*\right) .
\end{aligned}$$

We define the invasion fitness $F_i$, a measure of the ability of either virus to invade the host population,

$$F_i = \frac{p_i}{d+d_i} \text{ with } i \in \{x, y\} .$$

The ratio $d/g$ measures the replacement rate of healthy hosts.

In solution **V.1**, none of the competing viruses survives and the host population $H^* = g/d$ is constant at equilibrium. Solutions **V.2** and **V.3** correspond to the survival of only one of the viruses infecting the population of hosts, thus entailing the extinction of the other one. Finally, the degenerate solution **V.4** does not determine a unique value for $X^*$, indicating coexistence of the two competing viruses.

### 1.1 Stability analysis

The conditions for stability of the solutions are given by the sign of the eigenvalues of the Jacobian matrix associated to the system of equations. When all eigenvalues are negative, the fixed point is stable. When one of the eigenvalues is zero and the rest are negative, the fixed point degenerates into a continuous set of quasi-stable solutions. The Jacobian matrix, $J$, is the matrix of all first-order partial derivatives of a function or set of functions, and in our system takes the form:

$$J = \begin{pmatrix} \frac{\partial K_1}{\partial H} & \frac{\partial K_1}{\partial X} & \frac{\partial K_1}{\partial Y} \\ \frac{\partial K_2}{\partial H} & \frac{\partial K_2}{\partial X} & \frac{\partial K_2}{\partial Y} \\ \frac{\partial K_3}{\partial H} & \frac{\partial K_3}{\partial X} & \frac{\partial K_3}{\partial Y} \end{pmatrix} = \begin{pmatrix} -d - p_xX^* - p_yY^* & -p_xH^* & -p_yH^* \\ p_xX^* & p_xH^* - (d+d_x) & 0 \\ p_yY^* & p_yH^* - (d+d_y) & 0 \end{pmatrix} \tag{4}$$

where the functions $K_{1,2,3}$ are:

$$\begin{aligned}
K_1 &= g - dH - p_xXH - p_yYH \\
K_2 &= p_xXH - (d+d_x)X \\
K_3 &= p_yYH - (d+d_y)Y .
\end{aligned}$$

We evaluate the $J$ matrix of the system, Eq. ( 4) for every fixed point given by solutions **V.1** to **V.4** and find in explicit form, whenever possible, the roots or eigenvalues, $\lambda_i$, of the characteristic polynomial $p(\lambda) = 0$, where $p(\lambda) = det(J(H^*, X^*, Y^*) - \lambda\mathbb{I})$.

As an example, let us make explicit the calculation for solution **V.1**. First, we can substitute the values of $H^* = g/d$, $X^* = Y^* = 0$ in the Jacobian matrix, and then calculate the determinant of $J - \lambda\mathbb{I}$ (or the characteristic polynomial),

$$p(\lambda) = \left| \begin{pmatrix} -d-\lambda & -p_x H^* & -p_y H^* \\ 0 & p_x H^* - (d+d_x) - \lambda & 0 \\ 0 & p_y H^* - (d+d_y) & -\lambda \end{pmatrix} \right| ,$$

which yields

$$p(\lambda) = \lambda(d+\lambda)\left(p_x \frac{g}{d} - (d+d_x) - \lambda\right) .$$

There are three solutions for $p(\lambda) = 0$, $\lambda_1 = 0$, $\lambda_2 = -d$, and $\lambda_3 = p_x g/d - (d+d_x)$. The first eigenvalue is zero and the second one is always negative, since $d$ is by definition larger than zero. The condition for the third eigenvalue to be negative (and thus the solution stable) is obtained by requiring $\lambda_3 < 0$, that is, $F_x < d/g$, where we have used the definition of the invasion fitness above. Since $x$ and $y$ play interchangeable roles in model V, the solution has to be stable as well if Eqs. (2) and (3) are mutually swapped, so $F_y < d/g$ for full stability. The calculation proceeds analogously for all other solutions, occasionally involving some extra, though simple, algebra, to yield the following conditions:

- **V.1.** Conditions for stability, existence and non-negativity of $(H^*, X^*, Y^*) = \left(\frac{g}{d}, 0, 0\right)$ are
$$F_x < d/g \ \text{ and } \ F_y < d/g \, .$$
The conditions above show that viral invasions are precluded for a too slow growth of healthy hosts.
- **V.2.** Conditions for stability, existence and non-negativity of $(H^*, X^*, Y^*) = \left(\frac{d+d_x}{p_x}, \frac{g}{d+d_x} - \frac{d}{p_x}, 0\right)$ are
$$F_x > d/g \ \text{ and } \ F_x > F_y \, .$$
A virus with an invasion fitness above that of its competitor and simultaneously higher than the turnover of healthy hosts invades the population.
- **V.3.** Conditions for stability, existence and non-negativity of $(H^*, X^*, Y^*) = \left(\frac{d+d_y}{p_y}, 0, \frac{g}{d+d_y} - \frac{d}{p_y}\right)$ are:
$$F_y > d/g \ \text{ and } \ F_y > F_x \, .$$
- **V.4.** Conditions for stability, existence and non-negativity of $(H^*, X^*, Y^*) = \left(\frac{d+d_x}{p_x}, X^*, \frac{g}{d+d_y} - \frac{d}{p_y} - \frac{p_x}{p_y} X^*\right)$ are:
$$F_x = F_y > \frac{d}{g} \ \text{ and } \ \frac{g}{d+d_x} - \frac{d}{p_x} \ > \ X^* > 0 \, .$$
The conditions for coexistence of both viral types in this model are very stringent regarding parameter values, since only when their invasion fitness are equal and larger than the replacement of healthy hosts is stable coexistence possible.

### 2 ANALYTICAL SOLUTIONS AND STABILITY ANALYSIS OF MODEL S

Fixed points $(H^*, X^*, Y^*, S^*)$ of model S are the solutions of the system of equations

$$
\begin{aligned}
0 &= g - dH^* - p_x X^* H^* - p_y Y^* H^* - p_{sy} S^* H^* - p_{-y} S^* H^* \\
0 &= p_x X^* H^* - (d + d_x) X^* \\
0 &= p_y Y^* H^* - (d + d_y) Y^* - p_s Y^* S^* + p_{-y} S^* H^* \\
0 &= p_s Y^* S^* - (d + d_s) S^* + p_{sy} S^* H^* .
\end{aligned}
$$

As for model V, we only consider solutions that exist and are non-negative for all variables.

For $S^* = 0$ (no hosts infected with virus $y$ and satellite at equilibrium), four stable solutions are found:

$$
\textbf{S.1:}\ (H^*, X^*, Y^*, S^*) = \left(\frac{g}{d}, 0, 0, 0\right) \tag{5}
$$

$$
\textbf{S.2:}\ (H^*, X^*, Y^*, S^*) = \left(\frac{d + d_x}{p_x}, \frac{g}{d + d_x} - \frac{d}{p_x}, 0, 0\right) \tag{6}
$$

$$
\textbf{S.3:}\ (H^*, X^*, Y^*, S^*) = \left(\frac{d + d_y}{p_y}, 0, \frac{g}{d + d_y} - \frac{d}{p_y}, 0\right) \tag{7}
$$

$$
\textbf{S.4:}\ (H^*, X^*, Y^*, S^*) = \left(\frac{d + d_x}{p_x}, X^*, \frac{g}{d + d_y} - \frac{d}{p_y} - \frac{p_x}{p_y} X^*, 0\right) \tag{8}
$$

For $S^* \neq 0$, there are three additional solutions:

**S.5:** Extinction of class $X$,

$$
H^* = \frac{2gp_s}{Q \pm \sqrt{Q^2 + 4gp_s\left(-dp_s - p_y p_{sy} + \frac{(d+d_y)(p_{sy}+p_{-y})}{d+d_s}\right)}} \tag{9}
$$

$$
X^* = 0 \tag{10}
$$

$$
Y^* = \frac{d + d_s}{p_s} - \frac{p_{sy}}{p_s} H^* \tag{11}
$$

$$
S^* = -\frac{(d + d_y - p_y H^*)(d + d_s - p_{sy} H^*)}{p_s (d + d_s - (p_{sy} + p_{-y}) H^*)} , \tag{12}
$$

where $Q = p_s(d + g) + p_y(d + d_s) - (d + d_y)(p_{sy} + p_{-y})$.

Co-existence of $Y$ and $S$ populations in **S.5** implies that, with $S^* > 0$, helper virus populations $Y^* > 0$ can still replicate without the assistance of the satellite.

**S.6:** Co-existence of all classes has two different solutions, a stable fixed point, solution **S.6.a:**

$$H^* = \frac{d+d_x}{p_x} \tag{13}$$

$$X^* = \frac{g}{d+d_x} - \frac{d}{p_x} - \frac{p_y}{p_x}Y^* - \frac{p_{sy}+p_{-y}}{p_x}S^* \tag{14}$$

$$Y^* = \frac{d+d_s}{p_s} - \frac{p_{sy}}{p_s}H^* \tag{15}$$

$$S^* = -\frac{(d+d_y-p_yH^*)(d+d_s-p_{sy}H^*)}{p_s(d+d_s-(p_{sy}+p_{-y})H^*)}\,, \tag{16}$$

and a degenerate solution of type $(H^*, X(S^*), Y^*, S^*)$, for a range of values of $S^* > 0$ such that $X^* > 0$, solution **S.6.b:**

$$H^* = \frac{d+d_x}{p_x} \tag{17}$$

$$X^* = \frac{g}{d+d_x} - \frac{d}{p_x} - \frac{p_{-y}(d+d_y)}{p_xp_s} + \frac{p_{sy}+p_{-y}}{p_x} \quad S^* \tag{18}$$

$$Y^* = \frac{p_{-y}}{p_s}\frac{d+d_x}{p_x} \tag{19}$$

$$S^* > 0\,, \tag{20}$$

where we have used the extended definition of invasion fitness, as introduced in the Main text,

$$F_i = \frac{p_i}{d+d_i} \text{ with } i \in \{x, y, sy, c\}\,,$$

where for convenience we define $p_c = p_{sy} + p_{-y}$ (with $d_c = d_{sy} = d_s$). By definition, $F_c > F_{sy}$, since independent transmission is more likely than simultaneous transmission of the two entities.

### 2.1 Stability analysis

As above, we will determine the conditions on the model parameters ensuring that solutions are positive or zero, and seek for conditions yielding negative or zero eigenvalues, to determine stability.

The Jacobian matrix of model S reads

$$\begin{pmatrix} -d-p_xX^*-p_yY^*-p_cS^* & -p_xH^* & -p_yH^* & -p_cH^* \\ p_xX^* & p_xH^* & 0 & 0 \\ p_yY^*+p_{-y}S^* & 0 & -d-d_y+p_yH^*-p_sS^* & p_{-y}H^*-p_sY^* \\ p_{sy}S^* & 0 & p_sS^* & -d-d_s+p_sY^*+p_{sy}H^* \end{pmatrix}.$$

We evaluate the Jacobian matrix for all solutions in Eqs. (5–20) and solve the equation $\det(J(H^*, X^*, Y^*, S^*) - \lambda\mathbb{I}) = 0$. The conditions of stability that satisfy that all $\lambda_i \leq 0$ in the corresponding case are the following:

- **S.1.** Conditions for stability, existence and non-negativity of $(H^*, X^*, Y^*, S^*) = \left(\frac{g}{d}, 0, 0, 0\right)$ are

$$F_x < \frac{d}{g}, \quad F_y < \frac{d}{g} \quad \text{and} \quad F_{sy} < \frac{d}{g}\,.$$

The last condition adds to those in the previous model for the analogous solution. Healthy hosts can escape from infection if the host turnover $d/g$ of healthy hosts is above the infection fitness of any viruses $F_x, F_y$ or of the combination of virus and satellite, $F_{sy}$, when they are jointly transmitted.

- **S.2.** Conditions for stability, existence and non-negativity of $(H^*, X^*, Y^*, S^*) = \left(\frac{d+d_x}{p_x}, \frac{g}{d+d_x} - \frac{d}{p_x}, 0, 0\right)$ are

$$F_x > \frac{d}{g}, \quad F_x > F_y \text{ and } F_x > F_{sy}\,.$$

$F_x > F_{sy}$ is needed for invasion of $X$ populations: $x$ virus must overcome not only its competitor virus, but also its association with the satellite.

- **S.3.** Conditions for stability, existence and non-negativity of $(H^*, X^*, Y^*, S^*) = \left(\frac{d+d_y}{p_y}, 0, \frac{g}{d+d_y} - \frac{d}{p_y}, 0\right)$ are

$$F_y > \frac{d}{g}, \quad F_y > F_x, \text{ and } F_y\,(1-\gamma) > F_{sy}$$
$$\text{with} \quad \gamma = \frac{p_s}{d+d_s}\left(\frac{g}{d+d_y} - \frac{d}{p_y}\right).$$

- **S.4.** Conditions for stability, existence and non-negativity of $(H^*, X^*, Y^*, S^*) = \left(\frac{d+d_x}{p_x}, X^*, \frac{g}{d+d_y} - \frac{d}{p_y} - \frac{p_x}{p_y}X^*, 0\right)$ are

$$\begin{aligned} F_x = F_y &> \frac{d}{g}, \quad F_x = F_y > F_{sy}, \\ \frac{g}{d+d_x} - \frac{d}{p_x} > X^* &> \frac{g}{d+d_x} - \frac{d}{p_x} - \frac{p_y(d+d_s) - p_{sy}(d+d_y)}{p_x p_s}, \\ \frac{d+d_s}{p_s} - \frac{p_{sy}}{p_s}\frac{d+d_x}{p_x} &> Y^* > 0\,. \end{aligned}$$

- **S.5.** Conditions for stability, existence and non-negativity of the solution in Eqs. (9–12) are

$$H^* < \frac{d+d_x}{p_x},$$
$$\text{(i)} \quad H^* < 1/F_y \quad \text{and} \quad H^* < 1/F_c, \qquad \text{or} \qquad \text{(ii)} \quad H^* > 1/F_y \quad \text{and} \quad 1/F_{sy} > H^* > 1/F_c\,.$$

The conditions (i) are compatible with the conditions of stability and existence of **S.2**, resulting in a bistable regime where $F_c > F_x > F_y$ and both solutions **S.2** and **S.5** are possible stable equilibrium states. Depending on the initial conditions, the system approaches one or another solution. The region of bistability is shown in Fig. 5. Furthermore, as $H^*_{S.2} = \frac{d+d_x}{p_x}$ and $H^*_{S.5} < \frac{d+d_x}{p_x}$ to be stable, in the bistability the amount of healthy hosts with solution **S.5** will be always lower than the amount of healthy hosts in the solution **S.2** (as can be observed in Fig. 5). Note that the lower boundary of the region of bistability in Fig. 5 can be calculated by equating the solution for $H^*$ in Eq. (9) to $(d+d_x)/p_x$

(as for this value solution **S.5** becomes unstable) and the upper boundary can be calculated equating $F_x = F_y$ (as for this value solution **S.2** disappears and the degenerate solution **S.4** takes its place).

- **S.6.a.** Stability conditions for this solution cannot be analytically derived in explicit form. Conditions of existence and non-negativity are

$$\text{(i)} \quad F_y < F_x < F_c \quad \text{and} \quad F_x > F_{sy}\,, \quad \text{or} \quad \text{(ii)} \quad F_y > F_x > F_c\,.$$

Also, in order to obtain $X^* > 0$ the growth of healthy hosts must be sufficiently fast to verify

$$g > \frac{d+d_x}{p_x}\big(d + p_y Y^* + (p_{sy} + p_{-y})S^*\big)\,,$$

from where we obtain

$$g > F_x^{-1}\Big(d + \frac{p_{sy}}{p_s}(F_{sy}^{-1} - F_x^{-1})\Big(1 - \frac{F_y^{-1} - F_x^{-1}}{F_c^{-1} - F_x^{-1}}\Big)\Big)\,.$$

Numerically, we found that the solution in Eqs. (13–16) is stable when $F_y > F_x > F_c$. This extended region corresponds to the areas of coexistence in Figure 6, implying that **S.6.a** is a relevant solution defined in an extended domain. Note that the upper boundary of the region of coexistence shown in Fig. 6 can be calculated equating Eq. (14) to zero because in that boundary $X^* = 0$, or by equating the solution for $H^*$ in Eq. (9) to $(d + d_x)/p_x$ (as for this value solution **S.5** becomes unstable). The lower boundary can be calculated equating $F_x = F_y$ (as for this value $S^* = 0$ and in consequence **S.6.a** disappears and the degenerate solution **S.4** takes its place).

- **S.6.b.** Conditions for existence, non-negativity and stability of the degenerate solution in Eq. (20) are

$$F_x = F_y = F_c > \frac{d}{g}\,.$$

This is a highly degenerate solution that requires the coincidence of three invasion fitness, thus limiting its actual relevance.
